# Chromosome-scale inference of hybrid speciation and admixture with convolutional neural networks

**DOI:** 10.1101/2020.06.29.159673

**Authors:** Paul D. Blischak, Michael S. Barker, Ryan N. Gutenkunst

## Abstract

Inferring the frequency and mode of hybridization among closely related organisms is an important step for understanding the process of speciation and can help to uncover reticulated patterns of phylogeny more generally. Phylogenomic methods to test for the presence of hybridization come in many varieties and typically operate by leveraging expected patterns of genealogical discordance in the absence of hybridization. An important assumption made by these tests is that the data (genes or SNPs) are independent given the species tree. However, when the data are closely linked, it is especially important to consider their non-independence. Recently, deep learning techniques such as convolutional neural networks (CNNs) have been used to perform population genetic inferences with linked SNPs coded as binary images. Here we use CNNs for selecting among candidate hybridization scenarios using the tree topology (((*P*_1_, *P*_2_), *P*_3_), *Out*) and a matrix of pairwise nucleotide divergence (*d*_*XY*_) calculated in windows across the genome. Using coalescent simulations to train and independently test a neural network showed that our method, HyDe-CNN, was able to accurately perform model selection for hybridization scenarios across a wide-breath of parameter space. We then used HyDe-CNN to test models of admixture in *Heliconius* butterflies, as well as comparing it to a random forest classifier trained on introgression-based statistics. Given the flexibility of our approach, the dropping cost of long-read sequencing, and the continued improvement of CNN architectures, we anticipate that inferences of hybridization using deep learning methods like ours will help researchers to better understand patterns of admixture in their study organisms.

## INTRODUCTION

Hybridization is a widely observed phenomenon in nature that continues to be appreciated across diverse lineages in the Tree of Life. From some of the earliest systematic investigations of hybridization and introgression in irises (Anderson 1949; Anderson & Stebbins 1954) to more recent studies of selection on introgressed loci from Neanderthals into non-African humans (Harris & Nielsen 2016; Juric *et al.* 2016), understanding the importance of genetic exchanges between otherwise isolated lineages is of great interest to researchers across many fields in evolutionary genetics. Nevertheless, many studies aimed at inferring patterns of hybridization, especially in non-model taxa, lack the resolution to determine the specific mode of hybridization occurring within their study organism(s) (ie, homoploid hybrid speciation versus a pulse of admixture after divergence). However, the dropping cost of long-read sequencing technologies such as Oxford Nanopore and Pacific Biosciences are quickly democratizing the acquisition of chromosome-scale reference genomes for non-model species (Amarasinghe *et al.* 2020). As more labs obtain these references, it will be especially important to explore methods of hybridization inference that can leverage the information encoded by the contiguous variants along the genome.

In the absence of a genome reference, myriad summary statistics have been described for the detection of introgression and hybrid speciation from unlinked, biallelic SNPs. Among the earliest of these statistics is *D* (Green *et al.* 2010), or the “ABBA-BABA” statistic, which tests for a statistically significant excess of ABBA or BABA site patterns for biallelic sites evolving along genealogies from the species tree (((*P*_1_, *P*_2_), *P*_3_), Out). In the absence of introgression, we expect only incomplete lineage sorting (ILS) to generate ABBA or BABA site patterns, a result that derives from the coalescent model (Kingman 1982). Deviations from the coalescent expectations of site patterns evolving along genealogies forms the basis of nearly all tests for introgression, including tests on five-taxon trees (Eaton & Ree 2013; Pease & Hahn 2015), tests on *N*-taxon trees (Elworth *et al.* 2018), tests that assume a specific mode of hybridization (e.g., hybrid speciation; Blischak *et al.* 2018; Kubatko & Chifman 2019), as well as more recent approaches that incorporate sequence divergence into their test statistics (Hahn & Hibbins 2019; Hibbins & Hahn 2019; Forsythe *et al*. 2020). And while some of these statistics aim to distinguish among modes of hybridization and admixture or the direction of introgression, they all assume that genealogies across sites or genes are unlinked. The shared genealogical information contained in linked SNPs, especially about the size and coalescence time of linkage blocks, is an important feature of chromosome-scale data that is ignored in many inferences of hybridization, but which can likely provide greater power for distinguishing among more complex models.

Incorporating linkage into inferences of hybridization requires modeling both the coalescent process as well as the history of recombination along a chromosome, generating a highly complex object called the ancestral recombination graph (ARG; Hudson 1983; Griffiths & Marjoram 1996). However, because the complexity of the ARG typically makes it unwieldy to work with and nearly impossible to estimate, researchers developed an approximation called the sequentially Markovian coalescent (SMC; McVean & Cardin 2005), which greatly reduces the headaches associated with ARGs and creates an approximation that is both accurate and computationally tractable (but see methods Relate and tsinfer below). Applications of the SMC model have primarily come in the form of coalescent hidden Markov models (HMMs), which model genealogies or coalescent times (hidden states) of variants along the genome (observed states) such that adjacent variants either share a hidden state (no recombination) or are in different hidden states because of a recombination event (Hobolth *et al.* 2007; Dutheil *et al.* 2009). Some of the most well-known uses of coalescent HMMs are for the inference of pairwise coalescent times and population size trajectories. This includes methods that require phased genomes such as PSMC (Li & Durbin 2011) and MSMC (Schiffels & Durbin 2014), as well as methods that do not require known phase among the sampled individuals such as SMC++ (Terhorst *et al.* 2017). Two recent approaches, Relate (Speidel *et al.* 2019) and tsinfer (Kelleher *et al.* 2019), have even managed to use HMMs for approximating the full ARG by inferring the genealogical relationships among thousands of samples from entire chromosomes. Phylogenetic applications of coalescent HMMs include the original CoalHMM implementation for inferring parameters among human, chimp, and gorilla (Hobolth *et al.* 2007; Dutheil *et al.* 2009), although these approaches use a much simpler representation of the ARG that does not require the SMC model. Phylogenetic HMMs for the inference of admixture and hybridization have also been developed and include approaches such as PhyloNet-HMM (Liu *et al.* 2014) as well as updated version of the CoalHMM framework to model gene flow (Mailund *et al.* 2012).

The ability of coalescent HMMs and the SMC to infer complicated population histories comes from leveraging the correlation between genealogies along the chromosome resulting from linkage. However, as models become more complex, trying to incorporate and estimate all relevant parameters in a likelihood-based framework can become computationally prohibitive. Fortunately, efficient tools for simulating genome-scale data (e.g., msprime and SLiM; Kelleher *et al.* 2016; Haller & Messer 2019) provide opportunities to explore likelihood-free inference methods such as approximate Bayesian computation (ABC; Beaumont *et al.* 2002) and supervised machine learning (Sheehan & Song 2016; Schrider & Kern 2018). One particular type of supervised machine learning that has received increasing attention in population genomics and phylogenetics is deep learning, or the use of neural networks for model selection and/or parameter inference (LeCun *et al.* 2015). One type of deep learning algorithm that has recently been used to account for linkage among variant sites for different inference problems in population genomics is convolutional neural networks (CNNs; Flagel *et al.* 2018). Convolutional neural networks are typically employed for image recognition tasks because they focus on modeling structure in images by allowing adjacent pixels to share parameters in the network (LeCun *et al.* 1998). In a population genomic or phylogenetic setting, the input image for a CNN is simply any representation of genetic variation along the chromosome (e.g., a sequence alignment), typically among samples from a population or across species (Flagel *et al*. 2018; Suvorov *et al.* 2019). Because sites that are close to one another are more likely to come from the same ancestral recombination block, patterns of variation will reflect this correlation, allowing a CNN to be trained on genome-wide patterns of variation using simulations.

Here we explore the use of CNNs to perform model selection for different modes of hybridization using chromosome-scale representations of genomic data among pairs of species. Our approach uses simulations to train a CNN with pairwise sequence divergence calculated in windows across the chromosomes, which captures the correlation generated by linkage among (unphased) variant sites. Training, validation, and testing on independent data allowed us to evaluate the prediction accuracy of our trained CNNs to distinguish among models of hybridization and admixture. We then compared the prediction accuracy of the trained CNNs to random forest algorithms trained on a set of introgression-specific summary statistics to see if accounting for linkage with the CNN resulted in better accuracy for model selection. Finally, we used our CNN approach to test among different hypotheses for the mode of admixture between populations of *Heliconius* butterflies. Overall, we show that image-based representations of chromosome-scale, phylogenomic data lend themselves well to predicting models of hybridization. As genome reference sequences continue to become available, we anticipate that approaches like ours will be increasingly useful for not only inferring hybridization but for testing other hypotheses about evolutionary patterns as well.

## MATERIALS AND METHODS

### Representing phylogenomic data as an image

Using a similar conceptual setup as coalescent HMMs, we want to represent phylogenomic data that encodes both coalescence times along the chromosome and phylogenetic relationships among the sampled species. These two axes, chromosomal and phylogenetic, can be thought of as structuring genomic data by encoding linkage among variant sites along the chromosome and common ancestry across the phylogeny, such that a two-dimensional representation can be treated as an image used to train a CNN (see example image in Figure 1A). For the phylogenetic axis, we consider the four-taxon phylogeny typically used for tests of introgression, (((*P*_1_, *P*_2_), *P*_3_), *Out*), and represent it by considering all population pairs ordered by increasing phylogenetic distance: *P*_1_ × *P*_2_, *P*_1_ × *P*_3_, *P*_2_ × *P*_3_, *P*_1_ × *Out, P*_2_ × *Out, P*_3_ × *Out*. For the chromosomal axis, we first divide the chromosome into *W* windows. Next, within each window, we calculate the average number of nucleotide differences for all sampled chromosomes between each pair of species (Nei’s genetic distance; *d*_*XY,w*_) for species pairs *X* × *Y* and window *w* ∈ {1, …, *W*} (Nei & Li 1979; Nei 1987). *If there are n*_*X*_ and *n*_*Y*_ chromosomes from species *X* and *Y*, respectively, this gives us a distribution of *n*_*X*_ × *n*_*Y*_ values of *d*_*XY,w*_ within each window, which we treat as a proxy for the distribution of pairwise coalescent times between chromosomes sampled from each species. The relationship between coalescence times and genetic variation under the infinite sites model is a classical result in population genetics (Watterson 1975; Tajima 1983), and although other statistics for measuring genetic differences between populations exist (e.g., *F*_*ST*_ ; Wright 1931, 1943), we use *d*_*XY*_ here because it is a simple measure of divergence between chromosomes.

**Figure 1:**
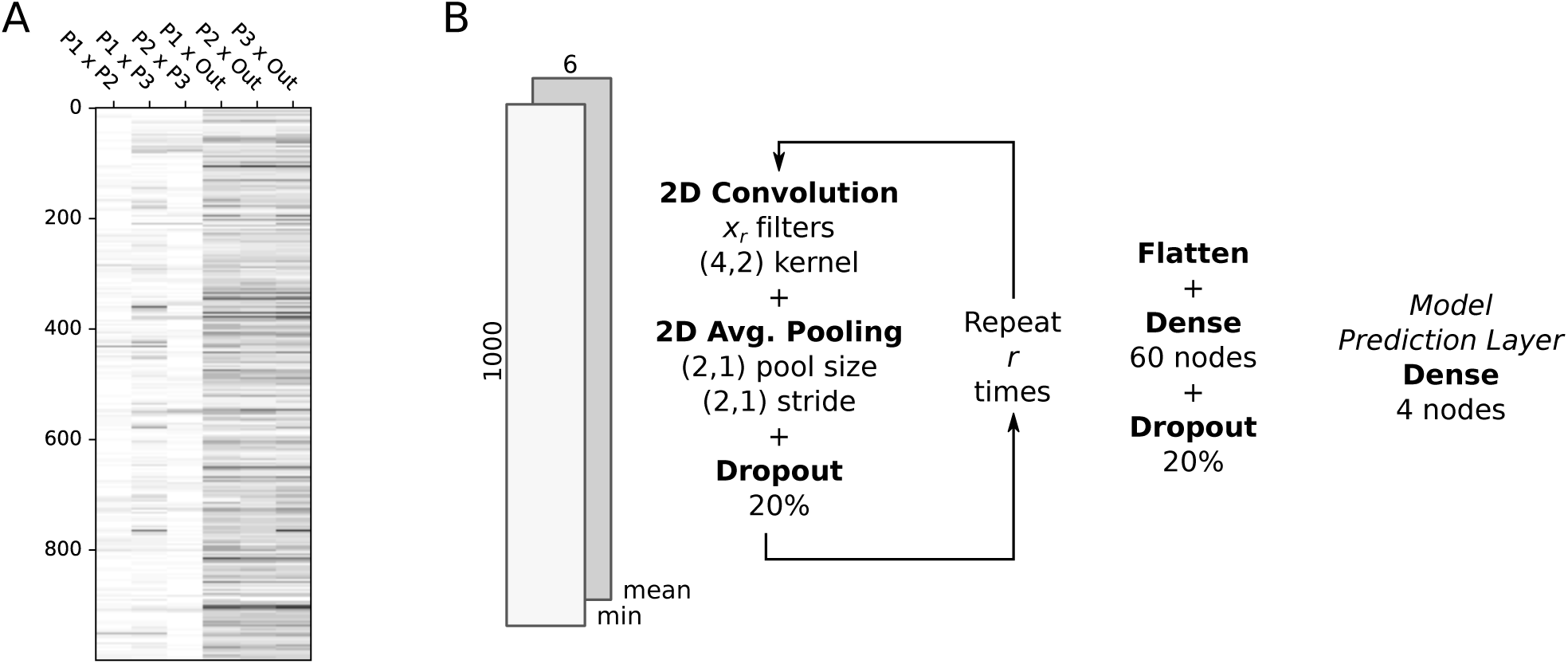
Neural network architecture for HyDe-CNN. (A) A sample image of minimum *d*_*XY*_ across 1 000 windows (rows) for the six pairwise comparisons among the species (columns). In this image, *P*_2_ is a hybrid species between *P*_1_ and *P*_3_, which is illustrated by the lower (lighter) values of *d*_*XY*_ for *P*_1_ × *P*_2_ and *P*_2_ × *P*_3_ compared to *P*_1_ × *P*_3_. (B) A simplified representation of the convolutional neural network architecture of HyDe-CNN.

### CNN architecture

The setup for our neural network is based on the LeNet architecture, which was introduced as one of the earliest uses of CNNs for the identification of hand-written characters (LeCun *et al*. 1998). In our implementation (Figure 1B), we start by repeating a sequence of three layers *r* times: a two-dimensional convolution layer with *x*_*r*_ filters, a two-dimensional average pooling layer, and a dropout layer. In our 2D convolution layer, a 4 × 2 filter walks across the input image, stepping across pixels in the rows and columns one at a time, and produces a single numerical output that is the convolution of the filter values (*f*) and the image values (*g*) at each step:

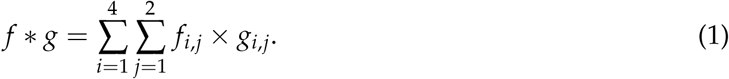

The values in the filter are randomly initialized such that the feature extraction from the underlying image data is stochastic. However, when the CNN weights are optimized during training, extracted features that lead to higher prediction accuracy will be given higher weight. The number of times this sequence of layers is repeated determines how reduced the original input image will be and therefore also determines the number of parameters that the network needs to estimate. For our network, we chose to repeat this sequence four times using filter sizes of 12, 24, 36, and 48. The next layer is a dense or fully connected layer with 60 nodes, followed by the final four-node dense layer that produces the model prediction. All layers except for the final layer use rectified linear unit activation functions (ReLU), which are constructed to try and capture nonlinear interactions in the input image (Agarap 2018). The final layer uses the softmax activation function, which produces output weights, *p*_*i*_ (∑_*i*_ *p*_*i*_ = 1), that are proportional to the support given to each model by the network. We refer to our architecture for hybridization inference as HyDe-CNN, which we specified with TensorFlow v2.1.0 using the tf.keras interface v2.2.4-tf (Abadi *et al.* 2016).

### Simulating training, validation, and test data

To test the ability of our data representation for distinguishing among models of hybridization with HyDe-CNN, we simulated chromosome-scale data under four models, depicted in Figure 2: no hybridization (1; *no_hyb*), hybrid speciation (2; *hyb_sp*), admixture (3; *admix*), and admixture with migration (4; *admix_mig*). All simulations were conducted in Python v3.7.6 with msprime v0.7.4 (Kelleher *et al.* 2016) using parameters drawn from the following distributions (numbers indicate which models use each parameter):

**Figure 2:**
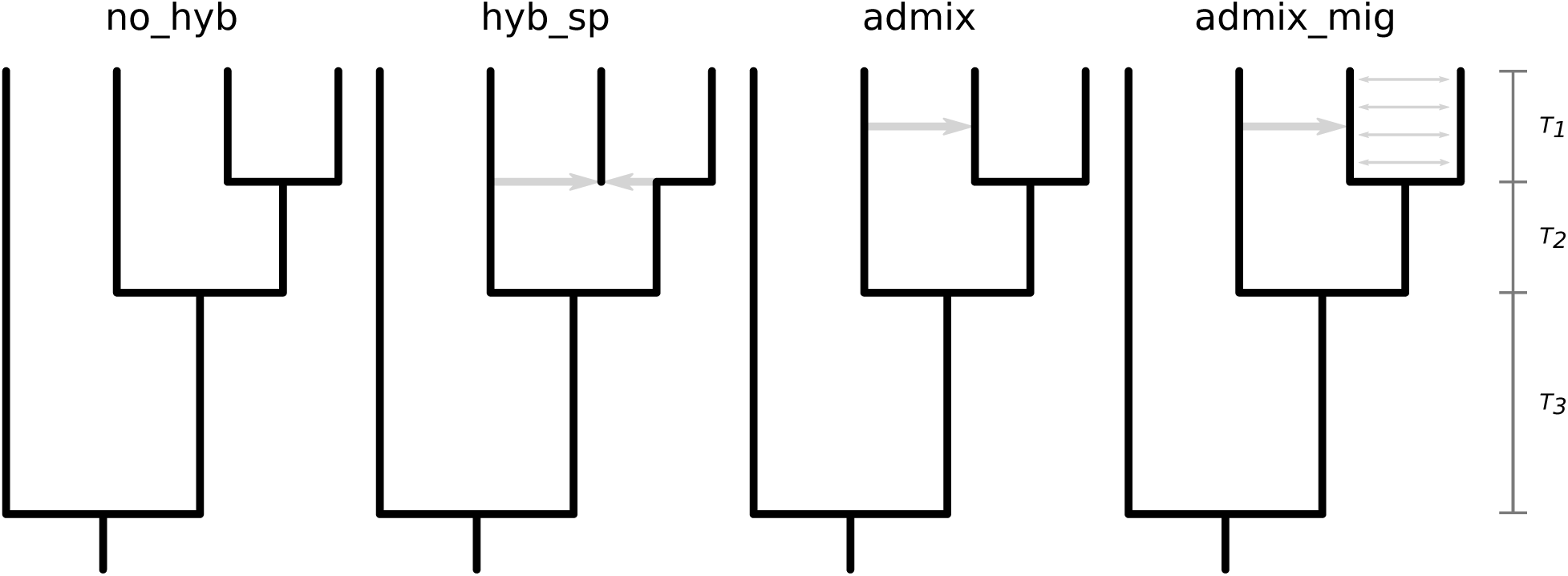
Models used to simulate genomic data for hybridization detection. From left to right they represent (1) no hybridization, (2) hybrid speciation, (3) pulsed admixture, and (4) pulsed admixture with ongoing gene flow. Parameters for all models are drawn from distributions as described in the **Materials and Methods**.

- [1–4] Sequence length (*L*) ∼ Discrete Uniform(1 × 10^7^, 5 × 10^7^, step = 1 × 10^5^).
- [1–4] Mutation rate (*µ*) and recombination rate (*r*) ∼ Uniform(2.5 × 10^*−*9^, 2.5 × 10^*−*8^).
- [1–4] Divergence times (*T*_1_, *T*_2_, *T*_3_) ∼ Gamma(10.0, 0.1), Gamma(20.0, 0.1), Gamma(40.0, 0.1).
- [3, 4] Admixture time (*T*_*mix*_) ∼ Uniform(0.1 × *T*_1_, 0.9 × *T*_1_).
- [2] Hybridization fraction (*γ*) ∼ Uniform(0.25, 0.75).
- [3, 4] Admixture proportion (*f*) ∼ Uniform(0.01, 0.25).
- [4] Migration rate (*m*) ∼ Uniform(2.5 × 10^*−*4^, 5 × 10^*−*4^).

Parameterizing the models in this way allowed us to explore a wide range of scenarios including variable hybridization and admixture times and proportions, variable mutation and recombination rates, and variable levels of sequence diversity (*θ* = 4*N*_*A*_ *Lµ* ranges from 100 to 5 000; *N*_*A*_ = 1 000). The resulting tree sequences from each msprime simulation run were summarized by calculating the number of pairwise differences between chromosomes for all pairs of species using 1 000 equally-spaced windows with the diversity() function in tskit v0.2.3 (Kelleher *et al.* 2018; Ralph *et al.* 2020). For each species, we sampled five chromosomes, leading to 25 pairwise values of *d*_*XY,w*_ within each window for each pair of species. To summarize this distribution of pairwise *d*_*XY,w*_, we calculated the minimum and mean values, storing these in numpy arrays for downstream CNN training, validation, and testing using the minimum alone, the mean alone, and the minimum and mean as two channels of the input image (Van Der Walt *et al.* 2011).

Next, we conducted 20 000 simulations for each model at three different branch scalings in coalescent units, 0.5 CUs (high ILS), 1.0 CUs (medium ILS), and 2.0 CUs (low ILS), for a total of 240 000 simulated data sets. In total we trained nine separate CNNs, one for each input type (minimum, mean, and minimum+mean) and branch scaling combination. For each CNN, we split the simulated data into training, validation, and test data using 15 000 images from each model for training and 2 500 images for both validation and testing. The input order of the images was shuffled so that images from different models were processed randomly by the CNN. Each individual image was also normalized by its maximum value to produce inputs with values between 0 and 1. Training and validation data were then fed to the CNN to fit the network parameters with a default learning rate of 0.001. We used the EarlyStopping callback within TensorFlow to stop training by monitoring the validation loss (CNN score on validation data) rather than setting a certain number of training iterations, which helps to prevent over-fitting to the training data. After training was completed, we ran the independent test samples through the final CNN to perform model selection. The model selection results were then processed with functions in scikit-learn v0.22.2 (Pedregosa *et al.* 2011) in Python to calculate prediction accuracy metrics and to generate confusion matrices.

To assess an alternative architecture to HyDe-CNN, we implemented a version of the network used by Flagel *et al.* (2018) for inferring introgression to train and test on our simulated input images. The biggest difference between HyDe-CNN and the network used by Flagel *et al*. is that it uses one-dimensional convolutional and average pooling layers, meaning that all pairs of species are used in these layers at once, rather than two at a time as in our architecture. Using one-dimensional layers in the network also restricted us to only test this architecture on the minimum and mean *d*_*XY*_ images since one-dimensional layers in TensorFlow cannot accept input data with more than one channel. All parameters in our implementation of the Flagel *et al*. architecture were kept the same except for the number of filters, which we reduced from 256 for the first layer and 128 for all other layers to 64 and 32, respectively. The primary reason for reducing the size of the network was because we trained all of our CNNs on a MacBook Pro laptop (2.3 GHz Intel Core i9, 16 GB RAM, 8 cores), which does not have a graphical processing unit capable of training a network of the original size specified by Flagel *et al*. All other training steps were completed exactly as above for the HyDe-CNN architecture.

### Assessing model accuracy

To further explore the predictions of the trained CNN, we conducted a second set of 10 000 simulations for each model and branch scaling combination to understand if prediction accuracy is associated with the actual biological parameters being used for the simulations. Data were simulated and processed into the formatted images of pairwise nucleotide divergence using the same methods described above. We focused here on the HyDe-CNN model trained using minimum pairwise divergence and processed all data sets in Python with TensorFlow. We then recorded the true model and parameter values used to simulate each data set, as well as the predicted model and its weight. Results were then plotted with GGPLOT2 v3.2.1 (Wickham 2009) in R v3.6.1 (R Core Team 2019) to visualize the relationship between the simulation parameters and model predictions.

### Comparison with summary statistics

For each of the 10 000 simulations from each model at the different branch scaling values used in the previous subsection, we also calculated a set of summary statistics constructed to detect introgression from biallelic SNP frequencies. We chose three statistics: *D* (Green *et al.* 2010), *f*_*hom*_ (Durand *et al.* 2011), *and D*_*p*_ (Hamlin *et al.* 2020). We then used the calculated values of the three statistics as predictor variables in a random forest classifier to jointly consider their ability to identify models of hybridization. To train a random forest classifier for each branch scaling, we split the simulated data sets from each model into 7 500 training and 2 500 testing samples and used the R packages ABCRF v1.8.1 (Pudlo et al. 2016) and CARET v6.0-86 (Kuhn 2008). Models were trained using the abcrf() function with 1 000 trees to build the random forest classifier and default values for all other options. The trained classifier was then used to predict the class of the testing samples and error rates were summarized using confusion matrices.

### Empirical example: Heliconius butterflies

Butterflies from the genus *Heliconius* have been studied extensively as a model system for non-bifurcating patterns of phylogenetic descent and the interplay of introgression and phenotypic evolution (Dasmahapatra *et al.* 2012; Nadeau *et al.* 2013; Edelman *et al.* 2019; Moest *et al.* 2020). Occurring primarily in Central and South America, *Heliconius* butterflies have conspicuous wing coloration patterning and are perhaps most famous for being one of the systems studied by Henry Walter Bates when describing his eponymous form of mimicry (Bates 1862). Previous work to uncover the extent of genetic exchanges among *Heliconius* species have quantified and described variation in amount of admixture between sympatric populations of different species (Martin *et al*. 2013, 2019). Here we sought to leverage this previous knowledge to further dissect these patterns of admixture using deep learning.

To test different models of admixture in *Heliconius*, we downloaded variant calls from Dryad for four taxa: *H. cydno* (*cyd*; *H. c. chioneus* + *H. c. zelinde*), *H. melpomene*-West (*mel*-W; *H. m. rosina* + *H. m. vulcanus*), *H. melpomene*-East (*mel*-E *H. m. malleti* + *H. m. amaryllis*), and *H. numata* (*num*) (https://doi.org/10.5061/dryad.sk2pd88; Martin *et al.* 2019). Previous work has shown that admixture occurs between sympatric *cyd* and *mel*-W west of the Andes mountains, typically going from *cyd* into *mel*-W (although some bidirectional introgression likely occured; Martin *et al.* 2019). We therefore wanted to test if we could distinguish between two alternative patterns of admixture between these two populations, (1) a single pulse of ancient admixture or (2) continuous gene flow at low frequencies, as well as including a model with no hybridization as a null hypothesis. To compare these models, we simulated data with msprime for 9.5 Mb of *Heliconius melpomene* chromosome five (positions 200 000 to 9 700 000) using a previously estimated recombination map for *Heliconius* (Davey *et al.* 2017) and the demographic model used for simulations in Martin et al. (2019). The fraction of admixture, *f*, was simulated using a random uniform distribution on the interval [0.3, 0.4], consistent with levels observed in these taxa Martin *et al.* (2013, 2019). The divergence time between *mel*-E and *mel*-W was 0.5 CU in the past, followed by the divergence between these two taxa and *cyd* at 1.5 CU, and finally the divergence of the outgroup, *num*, at 4.0 CU. To make the simulations more computationally feasible, the simulations were scaled to a population size of 2 000 individuals with a mutation rate of *µ* = 7.7 × 10^*−*7^. This mutation rate was chosen because it produced approximately the same number of segregating sites in the simulated data compared to the real data. For the ancient admixture model, we simulated the timing of admixture from a uniform distribution from [0.25, 0.5] CU in the past. For the continuous gene flow model, we divided the simulated value of *f* by the number of generations (0.5 CU × 2 × 2 000 = 1 000 generations) to give a cumulative level of admixture equal to *f* over the interval since the divergence of *mel*-W and *mel*-E.

We then simulated 20 000 data sets from each of the admixture models, plus the no hybridization scenario, using msprime and tskit as before to calculate *d*_*XY*_ among the population pairs in windows across the chromosome. We sampled 10 chromosomes for the populations representing *mel*-E, *mel*-W, and *cyd*, and sampled four chromosomes from the outgroup *num* population, dividing the 9.5 Mb of simulated sequence into 950 equally-sized, 10 kb windows. The distribution of pairwise *d*_*XY*_ values for each pair of populations was again calculated using the diversity() function and was summarized using the minimum, mean, and minimum+mean to generate three different types of input images for training. All aspects of training remained the same as for our original set of exploratory simulations, except for only having three models to choose from and using only 950 windows.

To test the predictions of the trained CNN for *Heliconius*, we used the pysam library (v0.15.3; https://github.com/pysam-developers/pysam) in Python to parse and process variant calls in VCF format on chromosome five using only positions 200 000 to 9 700 000 (Li *et al.* 2009). A total of 20 individuals (40 chromosomes) were sampled from each of *mel*-W, *mel*-E, and *cyd*, so instead of using all samples, we randomly drew five individuals (10 chromosomes) from each population to match our simulations. Only two individuals of *num* were available so we used both in all calculations. We repeated this random sampling of individuals 100 times, calculating pairwise values of *d*_*XY*_, summarizing the distribution using the minimum, mean, and minimum+mean, and running the resulting images through the corresponding trained CNN. We then assessed support for each of the models across the 100 replicates by recording how frequently each model received the highest weight.

## RESULTS

### Performance of trained CNNs

Training HyDe-CNN and the Flagel *et al*. network was not overly time consuming, usually taking between 5–10 minutes. It also usually only took a few training epochs for the validation loss score to plateau, suggesting that the networks quickly learn the information contained within the input images. This may also be because our networks are quite small compared to other more intensive implementations of deep neural networks (e.g., DeepVariant; Poplin *et al.* 2018), a point we return to in the **Discussion**. Prediction with the trained networks was only constrained by the amount of time it took to load the network into memory, making this step take just a few seconds. Not surprisingly, the most time intensive step for training the CNNs was generating the simulated input data images. Simulating 20 000 images from a single model usually took ∼24 hours but depended on the total number of generations needing to be simulated. However, because each simulated image coming from a model is independent, this step can by parallelized to speed up the generation of training data by splitting the simulation over many nodes on a computing cluster.

The trained HyDe-CNN architecture was able to achieve high accuracy for selecting among complex models of hybridization and admixture. Table 1 shows the overall prediction accuracy for both HyDe-CNN and the Flagel *et al*. network across input types and branch scaling factors. As expected, prediction accuracy increases as the branch scaling factor is increased, due to less conflicting signal in the data as a result of ILS. For both HyDe-CNN and the Flagel *et al*. architecture, the image encoding the minimum *d*_*XY*_ among pairs of species across the chromosome had the highest overall prediction accuracy across branch scaling factors. The two-channel minimum+mean CNN also had accuracy comparable to the CNN trained on just the minimum *d*_*XY*_. Interestingly, the CNNs trained on images of mean *d*_*XY*_ had much lower training and prediction accuracy regardless of architecture.

**Table 1:**
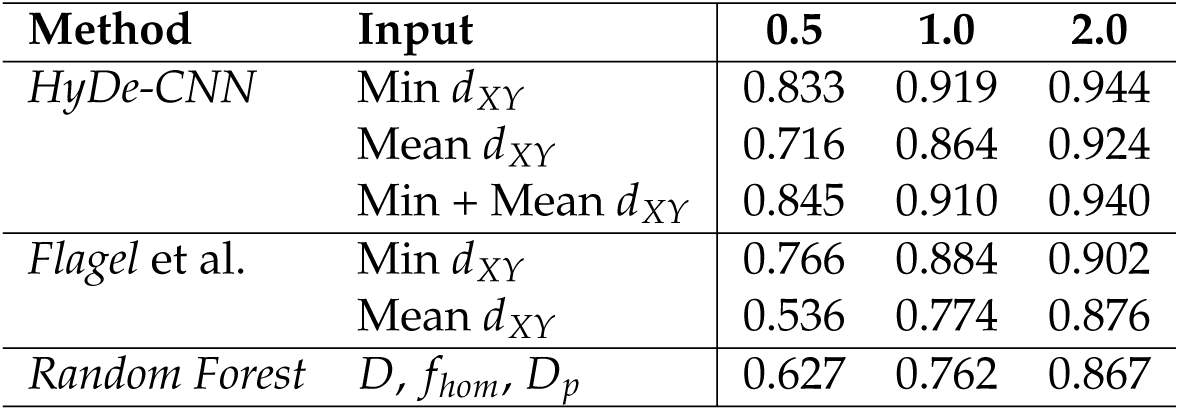
Overall prediction accuracy of the different machine learning algorithms used for model selection on independent test data. Branch scaling is given in coalescent units (CU).

| Method | Input | 0.5 | 1.0 | 2.0 |
| --- | --- | --- | --- | --- |
| *HyDe-CNN* | Min $d_{XY}$ | 0.833 | 0.919 | 0.944 |
| | Mean $d_{XY}$ | 0.716 | 0.864 | 0.924 |
| | Min + Mean $d_{XY}$ | 0.845 | 0.910 | 0.940 |
| *Flagel et al.* | Min $d_{XY}$ | 0.766 | 0.884 | 0.902 |
| | Mean $d_{XY}$ | 0.536 | 0.774 | 0.876 |
| *Random Forest* | $D, f_{hom}, D_p$ | 0.627 | 0.762 | 0.867 |

All trained models across both architectures primarily struggled to correctly identify the admixture model, especially at the 0.5 CU branch scaling. The confusion matrices in Figure 3 break this down for the HyDe-CNN minimum *d*_*XY*_ network, showing the proportion of correctly predicted models (diagonals) as well as how different models were incorrectly predicted (off-diagonals) across the different branch scalings. Precision and recall values for the HyDe-CNN minimum *d*_*XY*_ network are given in Table 2 and also show that false negatives (low recall) are primarily responsible for the low accuracy with the admixture model. The corresponding confusion matrix plots and precision/recall tables for the mean and minimum+mean HyDe-CNN predictions and the minimum and mean Flagel *et al*. network predictions are provided in the Supplemental Materials (Figures S1–S4 and Tables S1–S4).

**Table 2:** Precision and recall of HyDe-CNN trained on images of minimum *d*_*XY*_ for model selection on independent test data. Branch scaling is given in coalescent units (CU).

| Branch Scaling | Model | Precision | Recall |
| --- | --- | --- | --- |
| 0.5 | no_hyb | 0.841 | 0.924 |
|  | hyb_sp | 0.855 | 0.882 |
|  | admix | 0.777 | 0.640 |
|  | admix_mig | 0.846 | 0.885 |
| 1.0 | no_hyb | 0.928 | 0.965 |
|  | hyb_sp | 0.886 | 0.942 |
|  | admix | 0.919 | 0.774 |
|  | admix_mig | 0.942 | 0.995 |
| 2.0 | no_hyb | 0.945 | 0.980 |
|  | hyb_sp | 0.979 | 0.880 |
|  | admix | 0.874 | 0.916 |
|  | admix_mig | 0.983 | 1.000 |

**Figure 3:**
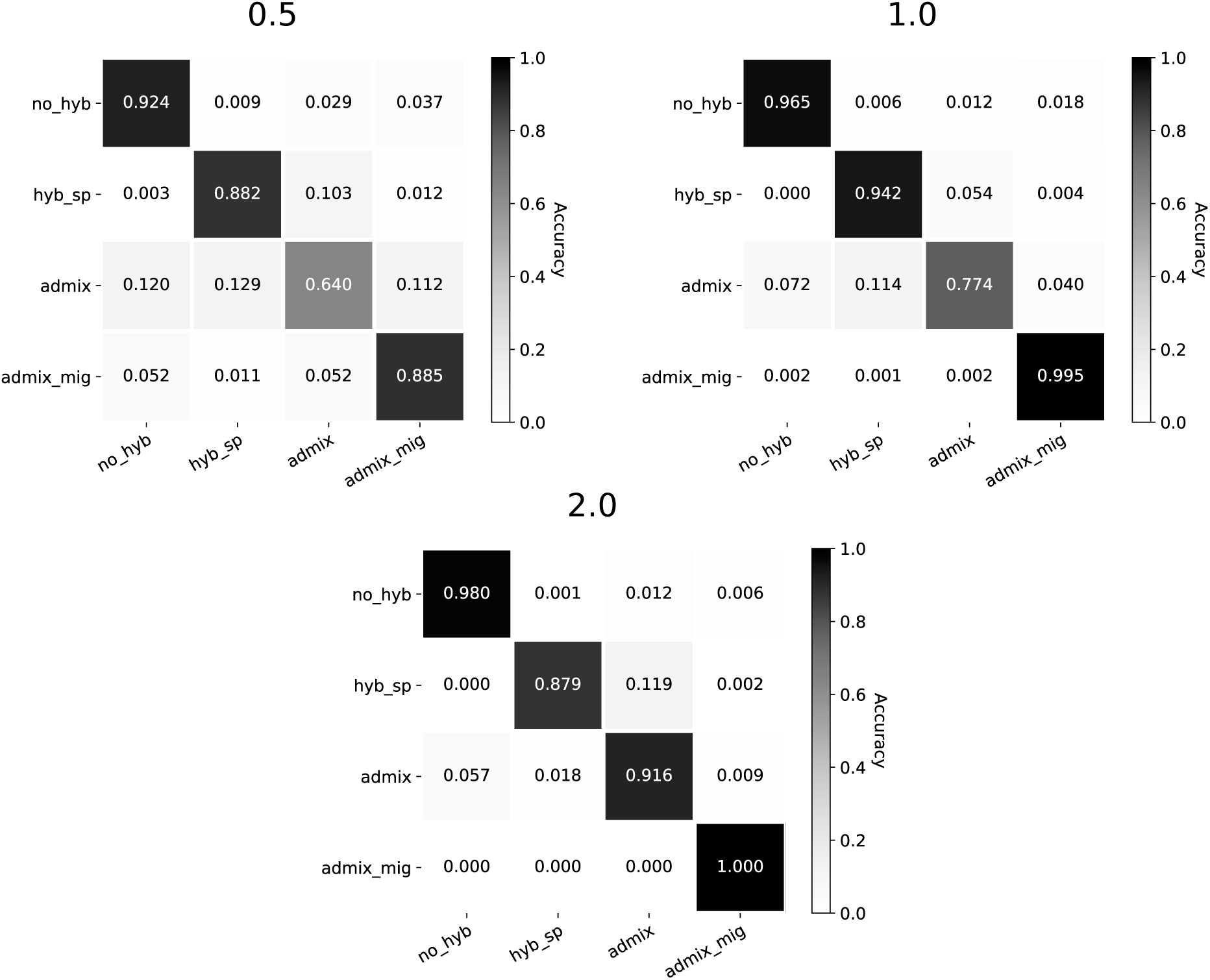
Confusion matrices for HyDe-CNN trained on minimum *d*_*XY*_. Branch scaling in coalescent units is noted above each plot. Model predictions were for 2,500 simulated images from each model. In each plot, rows correspond to the true model, columns correspond to the predicted model, and entries in the matrix show the proportion of simulations coming from each true model that are predicted to come from any one of the four possible models.

### Properties of misidentified models

Using 10 000 additional simulations from each model to explore how different parameters affect the prediction accuracy of our CNN, we found that models were typically misidentified in areas of parameter space where it would be biologically difficult to distinguish between the true model and the chosen model. We also found that when the best model was not the generating model, the CNN tended to give it lower weight, demonstrating more uncertainty in the model selection process. The mean weight for incorrect models ranged from 0.639–0.741, compared with the mean weight given when the true model was selected, which ranged from 0.906–0.944. The pattern of model misidentification based on the input parameters was most clearly seen with the admixture model, which was the model with the highest error rate across the majority of our tests. In Figure 4, we plot the predicted model for all 10 000 data sets simulated under the admixture model, as well as the two most important parameters distinguishing this model from the others: the admixture fraction (x-axis) and the timing of admixture (y-axis). As the coalescent branch lengths increase (top to bottom rows), we see that the majority of images are correctly predicted to come from the admixture model with high prediction weight from the CNN. When an image is misclassified, it is typically predicted to come from either the no hybridization or hybrid speciation model. However, this misclassification is not random. The no hybridization model is typically selected when the admixture fraction is close to 0. The hybrid speciation model is selected when the amount of admixture is higher (closer to 0.25) and when the timing of admixture is closer to the divergence between populations one and two. Similar plots for the no hybridization, hybrid speciation, and admixture with gene flow models are in the Supplemental Materials (Figures S5–S7).

**Figure 4:**
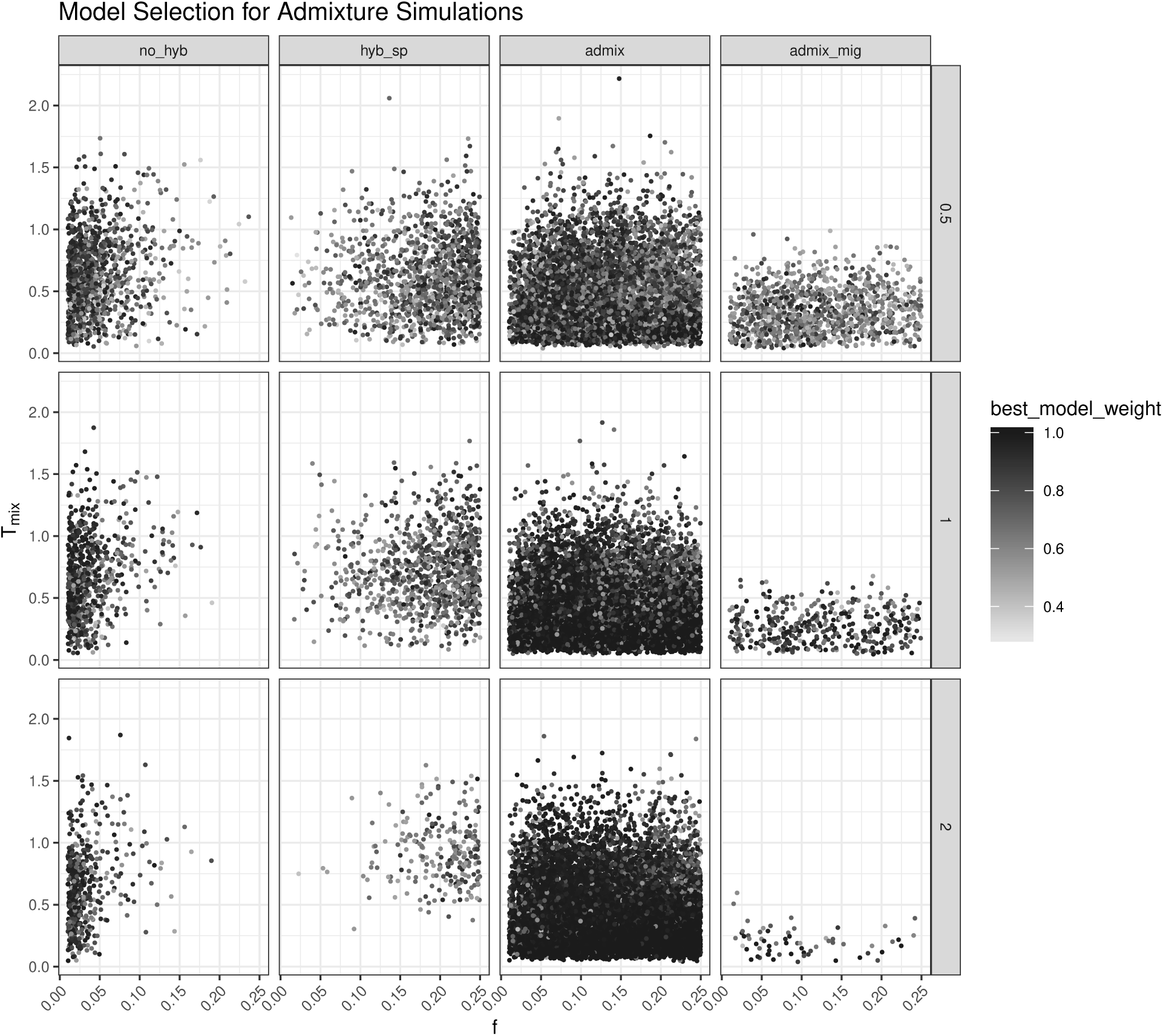
Scatterplot of model selection results for test simulations with the admixture model. Columns in the plot grid represent the predicted model and rows represent the different branch scaling factors in coalescent units. For each individual plot, the x-axis shows the fraction of admixture (*f*,), the y-axis shows the timing of admixture (*T*_*mix*_), and each dot represent the prediction for a single simulation (10 000 total). The shading of dots from light to dark corresponds to the predicted model weight (low to high, respectively).

### Comparison with summary statistics

The random forest classifier trained using summary statistics showed the same pattern of increasing training accuracy as branch scaling increased, having similar accuracy to the Flagel *et al*. network trained on minimum *d*_*XY*_ but lower accuracy when compared to HyDe-CNN across all input types (Table 1). The admixture model was again the most difficult to correctly classify, having a training misidentification rate ranging from 0.598 for the 0.5 CU simulations to 0.243 for the 2.0 CU simulations. Plotting the calculated test statistics across the models showed that there is considerable overlap in their values, which likely explains the poor prediction accuracy, especially at high levels of ILS (Figures S8–S10). Of the three statistics used, the variable importance metric reported by ABCRF showed that the *D*_*p*_ statistic was the most informative for performing classification. Precision and recall values for the reserved test data across branch scaling factors are given in Table S5 in the Supplemental Materials.

### Heliconius *butterflies*

Independent test data run through the trained HyDe-CNN architecture for *Heliconius* chromosome five showed that all input types (minimum, mean, minimum+mean) could distinguish among the three models with high accuracy (Figure S11). Using the trained networks to test for the mode of admixture between *mel*-W and *cyd* using 100 random samples of 5 individuals from each of the three ingroup populations (*mel*-E, *mel*-W, *cyd*) showed that the ancient admixture model was the predicted model for 98% of the minimum *d*_*XY*_ images, 100% of the mean *d*_*XY*_ images, and 100% of the minimum+mean images. The average model weight across the 100 runs for the admixture model was 0.885 for the minimum *d*_*XY*_ network, 0.988 for the mean *d*_*XY*_ network, and 0.991 for the minimum+mean *d*_*XY*_ network.

## DISCUSSION

Compared to traditional, introgression-specific summary statistics, as well as a previously employed CNN architecture (Flagel *et al.* 2018), our HyDe-CNN approach produced model predictions with the highest levels of accuracy, precision, and recall, especially at the shortest branch scaling (0.5 CU) where the signal for hybridization is expected to be the most noisy. In particular, the networks that included minimum *d*_*XY*_ performed the best for predicting patterns of admixture. This makes sense given that hybridization leads to coalescent events that are more recent than the divergence times between population pairs, which is more likely to be captured by the minimum coalescent time between sampled chromosomes. Inferences of species divergence times and hybridization based on minimum coalescent times (Kubatko 2009; Kubatko *et al.* 2009) or deviations from expected coalescent times (Joly *et al.* 2009) provided some of the first implementations of tests for hybridization. Using minimum *d*_*XY*_ within windows across the genome with HyDe-CNN therefore builds directly on these early ideas, providing a way to encode not only summaries of coalescent times but also their correlation across the genome due to linkage.

In our testing of HyDe-CNN, we deliberately chose to sample parameters from distributions rather than picking a small set of values to use for our simulations. In this way, we were able to test the robustness of the CNN’s predictions across a wide swath of parameter space, showing that our method is capable of accurately selecting among competing hybridization models. For tests on empirical data, the ability to incorporate uncertainty for certain parameters is especially appealing, particularly for organisms where not as much is known about their evolutionary history. Our *Heliconius* analysis illustrates this flexibility well as we put distributions on the timing and magnitude of admixture but used fixed values for the rest of the input parameters. Nevertheless, the trained network showed high power for selecting among the candidate models (Figure S11) and overwhelmingly supported the ancient admixture model when analyzing data from chromosome five, a result that agrees with previous work (Martin *et al.* 2019). Current likelihood-free approaches such as ABC also allow for incorporating uncertainty when estimating evolutionary parameters and performing model selection (Beaumont *et al.* 2002). However, obtaining estimates of parameters with ABC relies on sampling all parameters in proportion to their approximate posterior probability, which can require a huge amount of simulation (Beaumont 2010, the “curse of dimensionality”). Methods that combine machine learning and ABC to reduce the amount of needed simulation are also being developed (Pudlo *et al.* 2016; Estoup *et al.* 2018), and are currently being used for applications in model selection for species delimitation (Smith & Carstens 2020) and demographic inference (Smith *et al.* 2017). Incorporating uncertainty estimates in convolutional neural networks are also beginning to emerge (Laumann *et al.* 2018) and are a potentially promising direction for extending our approach.

Another desirable feature of our HyDe-CNN method is that the trained network failed to correctly classify models in ways that reflect our biological expectations. While it may seem odd to highlight our method’s failure, it is important to emphasize the interpretability of HyDe-CNN’s predictions based on the parameters of the simulations that are used to train it. The best example of this is the method’s difficulty with predicting the admixture model. The parameter bounds that we placed on this model for the timing and amount of admixture were chosen to approach the bounds of both the no hybridization and hybrid speciation models. With the no hybridization model, only admixture scenarios with low amounts of genetic exchange were incorrectly inferred to come from a model with no admixture (Figure 4; first column). Similarly, when admixture models were mistakenly inferred for the hybrid speciation model, it was when the amount of admixture was close to 25%, the lower bound for *γ* in the hybrid speciation model (Figure 4; third column). Failing to correctly select the admixture model in these cases, particularly when ILS is high (0.5 CU simulations), as well as selecting the most biologically similar model rather than a random model, suggests to us that HyDe-CNN is able to successfully model real biology. Furthermore, the fact that HyDe-CNN typically gives less weight to incorrectly selected models suggests that the trained network is able to capture some aspects of uncertainty in the model selection process. Future work to decode the impact of the intermediate layers of the network, as well as determining which parts of the input images are most informative for model selection, will hopefully help to further uncover how biology plays into the model selection process for CNNs.

Exploring more ways to extract information from genomic data using deep learning is likely to continue being an active area of research not only in terms of designing neural network architectures but also for thinking about how to represent genetic variation. The way that we represent genomic data as an image differs from other approaches that use the actual nucleotide alignment or genotype matrix as their input for CNN training (Flagel *et al.* 2018; Suvorov *et al.* 2019; Battey *et al.* 2020). We chose our data representation because we believed it to be an intuitive summary of pairwise coalescence times between species organized by the pattern of divergence in the underlying phylogeny. However, as we mentioned briefly above, our network architecture is both smaller and simpler than other CNNs used in population genetics. On the one hand, this makes the network easy to train, does not require the use of a GPU, and is less likely to lead to overfitting, all while still generalizing to making predictions on independent test data. On the other hand, it is possible that a larger network could provide more power to tease apart subtle differences between similar models (e.g., admixture versus hybrid speciation). More sophisticated and larger network architectures, such as exchangeable (Chan *et al.* 2018) and recurrent (Adrion *et al.* 2020) neural networks, have led to powerful inferences of recombination landscapes and could potentially help to better leverage the linkage information encoded by our representation. As more researchers begin working with deep learning models, we look forward to the continued testing and refinement of methods for not only estimating patterns of hybridization but for making other phylogenomic inferences as well.

## ACKNOWLEDGEMENTS

The authors thank M. L. Smith for the invitation to submit this paper for the special issue on Machine Learning in Molecular Ecology. We also thank J. E. James and S. Martin for help with processing and modeling the *Heliconius* data, as well as the *Heliconius* community in general for making their excellent genomic resources publicly available. In addition, we are grateful to members of the Barker and Gutenkunst labs for comments that helped to improve this manuscript. This work was supported by a National Science Foundation Postdoctoral Research Fellowship in Biology (IOS-1811784 to P.D.B.) and by the National Institute of General Medical Sciences of the National Institutes of Health (R01GM127348 to R.N.G.).

## DATA ACCESSIBILITY STATEMENT

Code for all simulations and analyses is available on GitHub (https://github.com/pblischak/hyde-cnn.git) with additional documentation on ReadTheDocs (https://hyde-cnn.readthedocs.io). All simulated input images and trained models are deposited on Dryad (DOI:####).

## AUTHOR CONTRIBUTIONS

PDB designed the study with guidance from MSB and RNG. PDB performed the simulations and data analyses. PDB wrote the initial manuscript draft with input from MSB and RNG. All authors read and approved the final manuscript.

