## Supplemental Materials for "Chromosome-scale inference of hybrid speciation and admixture with convolutional neural networks"

Paul D. Blischak, Michael S. Barker, and Ryan N. Gutenkunst

### Supplemental Text

#### S1 Summary statistics

To train a random forest classifier for model selection we calculated the statistics $D$ (Green *et al.* 2010), $f_{hom}$ (Durand *et al.* 2011), and $D_p$ (Hamlin *et al.* 2020) on the phylogeny $(((P_1, P_2), P_3), O)$. The allele-frequency based formulas for each statistic are given below, where $\hat{p}_{i,\ell}$ is the derived allele frequency for population $i$ at locus $\ell$.

$$D = \frac{\sum_L (1-\hat{p}_{1,\ell})\hat{p}_{2,\ell}\hat{p}_{3,\ell}(1-\hat{p}_{O,\ell}) - \hat{p}_{1,\ell}(1-\hat{p}_{2,\ell})\hat{p}_{3,\ell}(1-\hat{p}_{O,\ell})}{\sum_L (1-\hat{p}_{1,\ell})\hat{p}_{2,\ell}\hat{p}_{3,\ell}(1-\hat{p}_{O,\ell}) + \hat{p}_{1,\ell}(1-\hat{p}_{2,\ell})\hat{p}_{3,\ell}(1-\hat{p}_{O,\ell})}.$$

$$f_{hom} = \frac{\sum_L (1-\hat{p}_{1,\ell})\hat{p}_{2,\ell}\hat{p}_{3,\ell}(1-\hat{p}_{O,\ell}) - \hat{p}_{1,\ell}(1-\hat{p}_{2,\ell})\hat{p}_{3,\ell}(1-\hat{p}_{O,\ell})}{\sum_L (1-\hat{p}_{1,\ell})\hat{p}_{3,\ell}\hat{p}_{3,\ell}(1-\hat{p}_{O,\ell}) - \hat{p}_{1,\ell}(1-\hat{p}_{3,\ell})\hat{p}_{3,\ell}(1-\hat{p}_{O,\ell})}.$$

$$D_p = \frac{\sum_L (1-\hat{p}_{1,\ell})\hat{p}_{2,\ell}\hat{p}_{3,\ell}(1-\hat{p}_{O,\ell}) - \hat{p}_{1,\ell}(1-\hat{p}_{2,\ell})\hat{p}_{3,\ell}(1-\hat{p}_{O,\ell})}{\sum_L (1-\hat{p}_{1,\ell})\hat{p}_{2,\ell}\hat{p}_{3,\ell}(1-\hat{p}_{O,\ell}) + \hat{p}_{1,\ell}(1-\hat{p}_{2,\ell})\hat{p}_{3,\ell}(1-\hat{p}_{O,\ell}) + \hat{p}_{1,\ell}\hat{p}_{2,\ell}(1-\hat{p}_{3,\ell})(1-\hat{p}_{O,\ell})}.$$

### Supplemental Tables

Table S1: Precision and recall of HyDe-CNN trained on images of mean $d_{XY}$ for model selection on independent test data. Branch scaling is given in coalescent units (CU).

| Branch Scaling | Model | Precision | Recall |
|---|---|---|---|
| 0.5 | no_hyb | 0.725 | 0.850 |
| | hyb_sp | 0.796 | 0.896 |
| | admix | 0.572 | 0.365 |
| | admix_mig | 0.708 | 0.753 |
| 1.0 | no_hyb | 0.827 | 0.960 |
| | hyb_sp | 0.902 | 0.908 |
| | admix | 0.804 | 0.700 |
| | admix_mig | 0.923 | 0.889 |
| 2.0 | no_hyb | 0.908 | 0.989 |
| | hyb_sp | 0.945 | 0.914 |
| | admix | 0.901 | 0.803 |
| | admix_mig | 0.943 | 0.992 |

Table S2: Precision and recall of HyDe-CNN trained on images of minimum+mean $d_{XY}$ for model selection on independent test data. Branch scaling is given in coalescent units (CU).

| Branch Scaling | Model | Precision | Recall |
|---|---|---|---|
| 0.5 | no_hyb | 0.837 | 0.930 |
| | hyb_sp | 0.926 | 0.841 |
| | admix | 0.798 | 0.674 |
| | admix_mig | 0.822 | 0.932 |
| 1.0 | no_hyb | 0.911 | 0.972 |
| | hyb_sp | 0.843 | 0.982 |
| | admix | 0.971 | 0.691 |
| | admix_mig | 0.944 | 0.996 |
| 2.0 | no_hyb | 0.916 | 0.985 |
| | hyb_sp | 0.930 | 0.949 |
| | admix | 0.943 | 0.826 |
| | admix_mig | 0.972 | 1.000 |

Table S3: Precision and recall of the Flagel *et al.* network trained on images of minimum $d_{XY}$ for model selection on independent test data. Branch scaling is given in coalescent units (CU).

| Branch Scaling | Model | Precision | Recall |
|---|---|---|---|
| 0.5 | no_hyb | 0.810 | 0.808 |
| | hyb_sp | 0.859 | 0.845 |
| | admix | 0.703 | 0.457 |
| | admix_mig | 0.699 | 0.956 |
| 1.0 | no_hyb | 0.855 | 0.975 |
| | hyb_sp | 0.922 | 0.858 |
| | admix | 0.789 | 0.770 |
| | admix_mig | 0.979 | 0.932 |
| 2.0 | no_hyb | 0.899 | 0.936 |
| | hyb_sp | 0.956 | 0.817 |
| | admix | 0.776 | 0.856 |
| | admix_mig | 0.996 | 0.997 |

Table S4: Precision and recall of the Flagel *et al.* network trained on images of mean $d_{XY}$ for model selection on independent test data. Branch scaling is given in coalescent units (CU).

| Branch Scaling | Model | Precision | Recall |
|---|---|---|---|
| 0.5 | no_hyb | 0.398 | 0.668 |
| | hyb_sp | 0.907 | 0.738 |
| | admix | 0.494 | 0.474 |
| | admix_mig | 0.483 | 0.263 |
| 1.0 | no_hyb | 0.671 | 0.846 |
| | hyb_sp | 0.948 | 0.792 |
| | admix | 0.664 | 0.496 |
| | admix_mig | 0.833 | 0.964 |
| 2.0 | no_hyb | 0.853 | 0.941 |
| | hyb_sp | 0.905 | 0.894 |
| | admix | 0.837 | 0.701 |
| | admix_mig | 0.905 | 0.970 |

Table S5: Precision and recall of the random forest classifier trained with introgression summary statistics for model selection on independent test data. Branch scaling is given in coalescent units (CU).

| Branch Scaling | Model | Precision | Recall |
|---|---|---|---|
| 0.5 | no_hyb | 0.651 | 0.765 |
| | hyb_sp | 0.812 | 0.813 |
| | admix | 0.471 | 0.381 |
| | admix_mig | 0.541 | 0.549 |
| 1.0 | no_hyb | 0.784 | 0.867 |
| | hyb_sp | 0.838 | 0.836 |
| | admix | 0.665 | 0.560 |
| | admix_mig | 0.744 | 0.784 |
| 2.0 | no_hyb | 0.877 | 0.944 |
| | hyb_sp | 0.849 | 0.852 |
| | admix | 0.804 | 0.755 |
| | admix_mig | 0.936 | 0.917 |

### Supplemental Figures

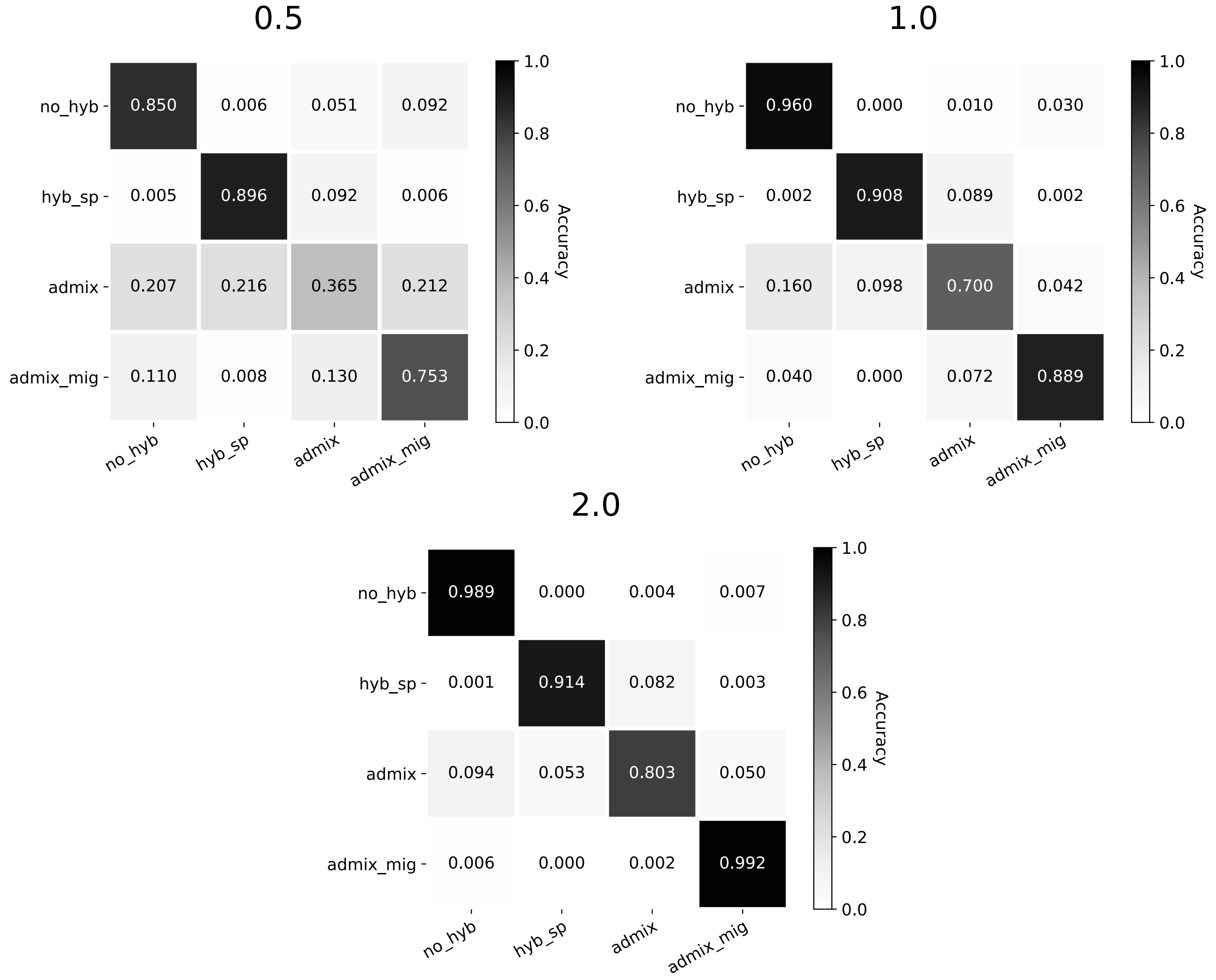

Figure S1: Confusion matrices for the HyDe-CNN architecture trained on images of mean $d_{XY}$ across different branch scaling factors (specified in coalescent units).

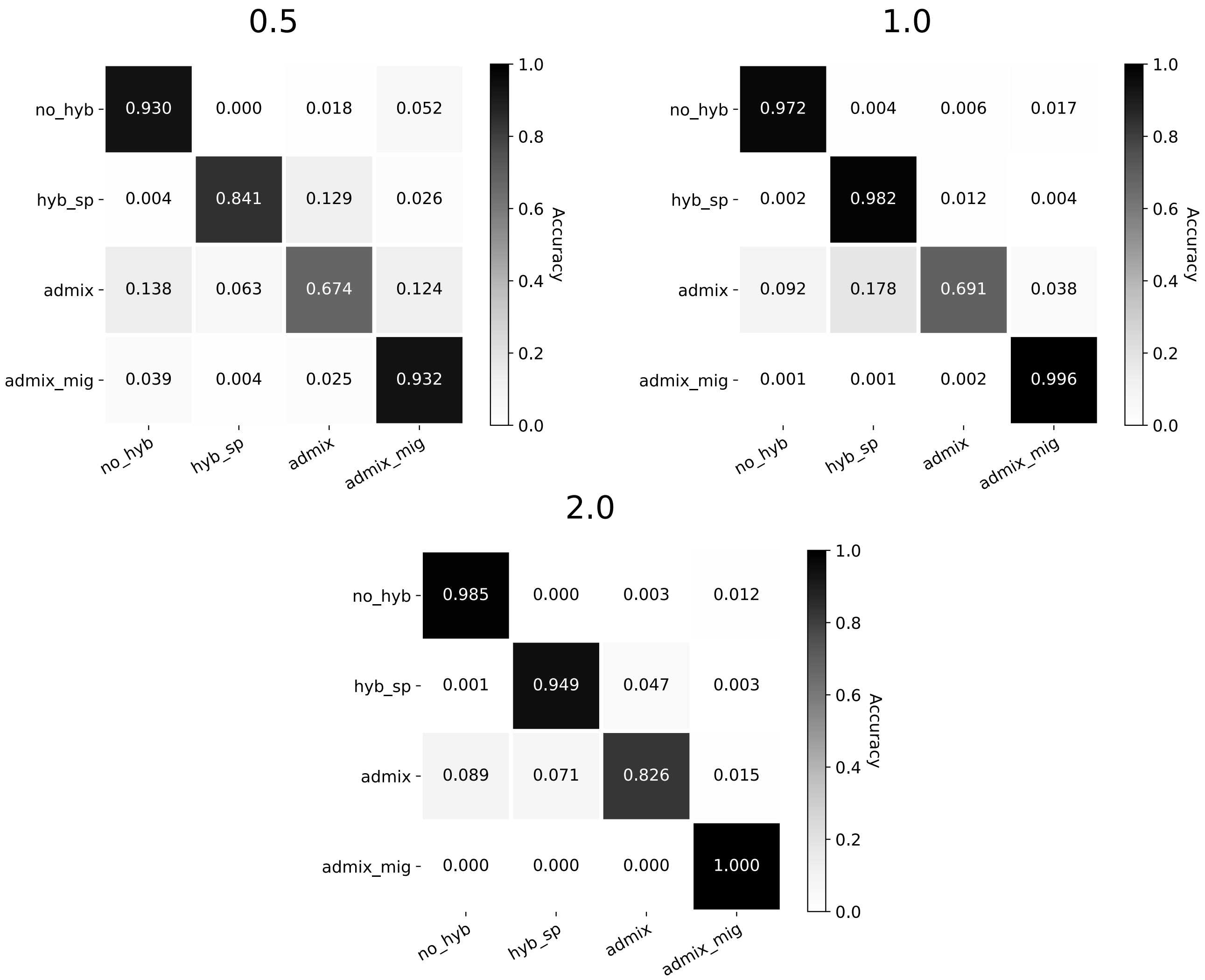

Figure S2: Confusion matrices for the HyDe-CNN architecture trained on images of minimum+mean $d_{XY}$ across different branch scaling factors (specified in coalescent units).

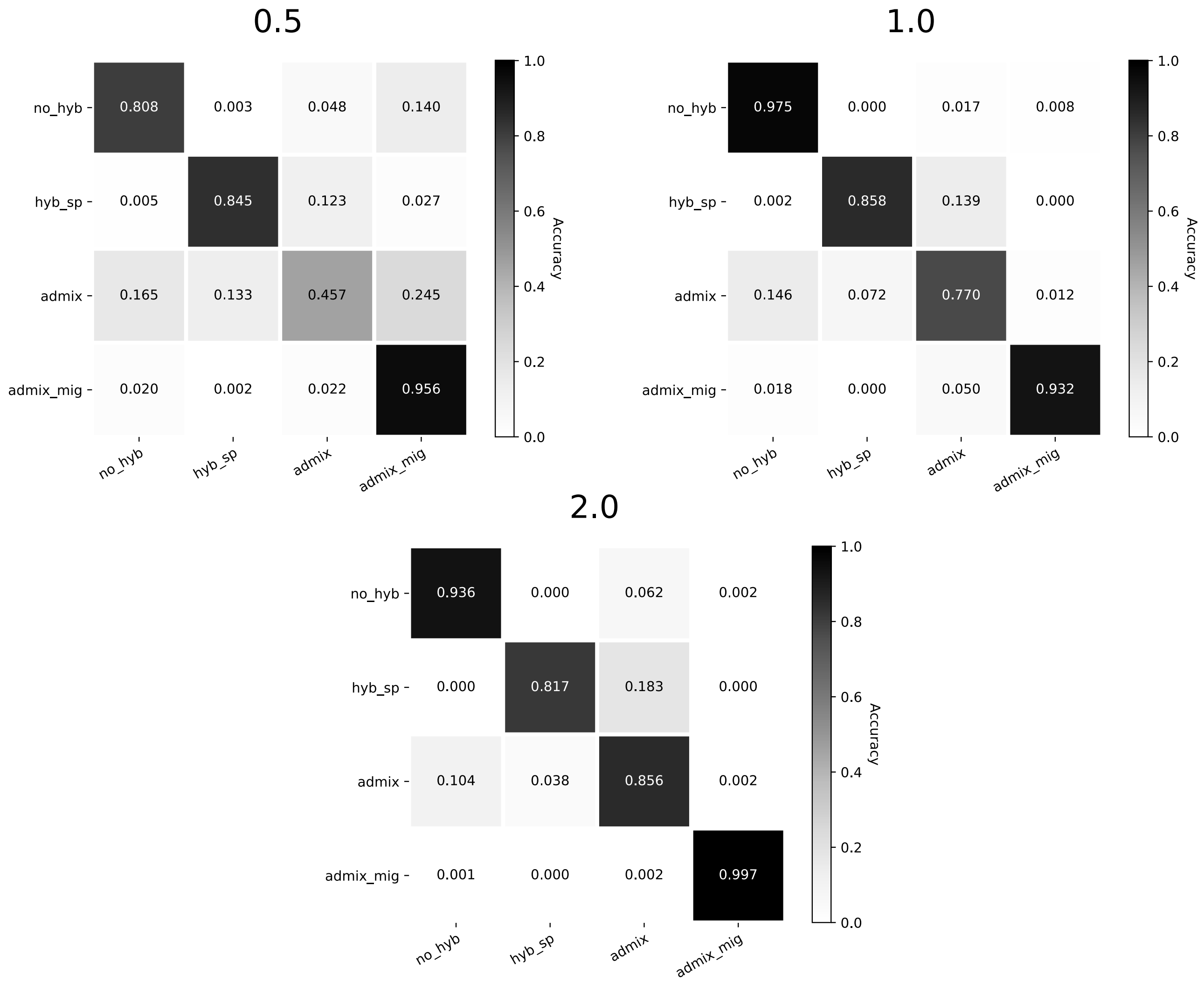

Figure S3: Confusion matrices for the Flagel *et al.* network trained on images of minimum $d_{XY}$ across different branch scaling factors (specified in coalescent units).

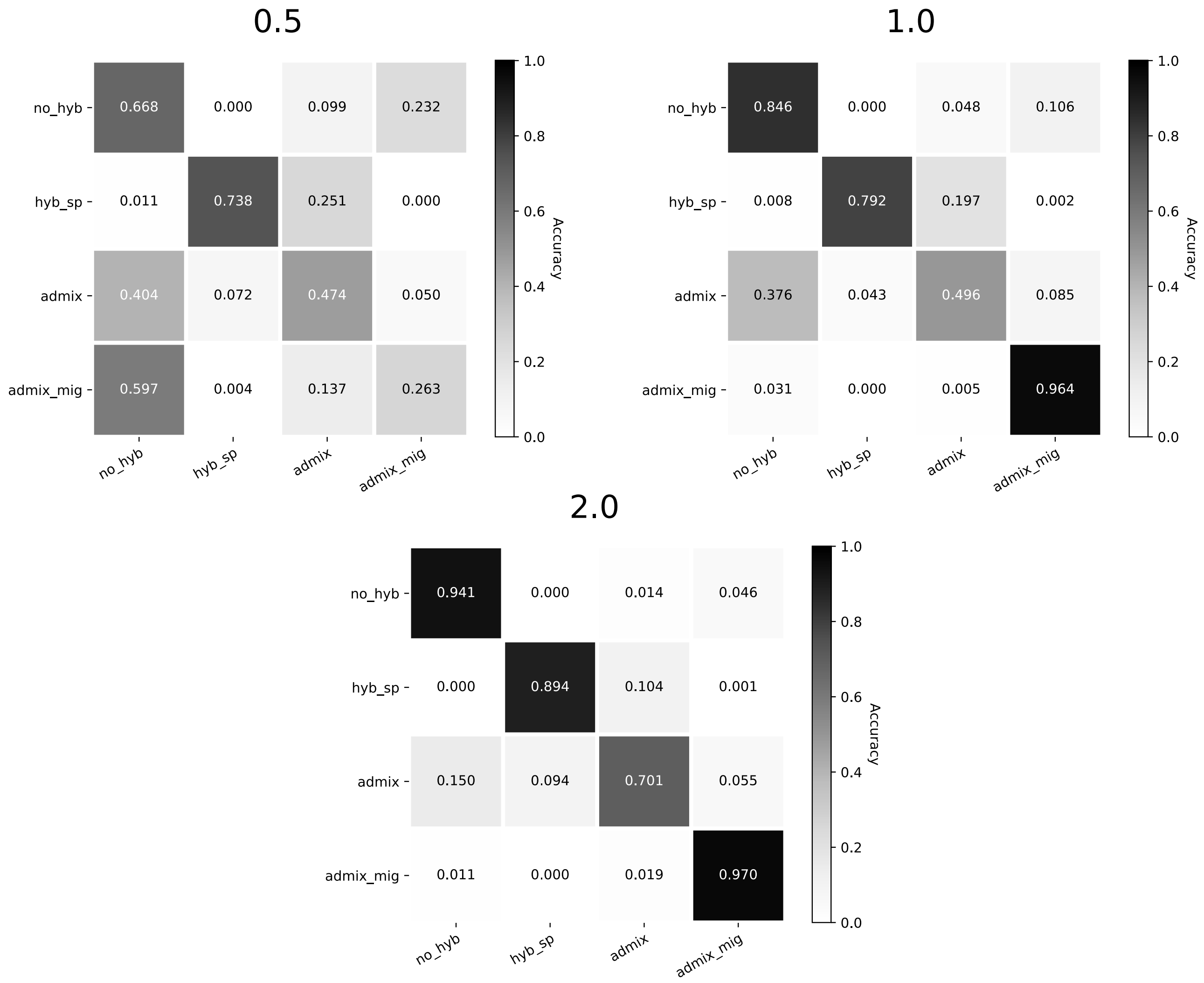

Figure S4: Confusion matrices for the Flagel *et al.* network trained on images of mean $d_{XY}$ across different branch scaling factors (specified in coalescent units).

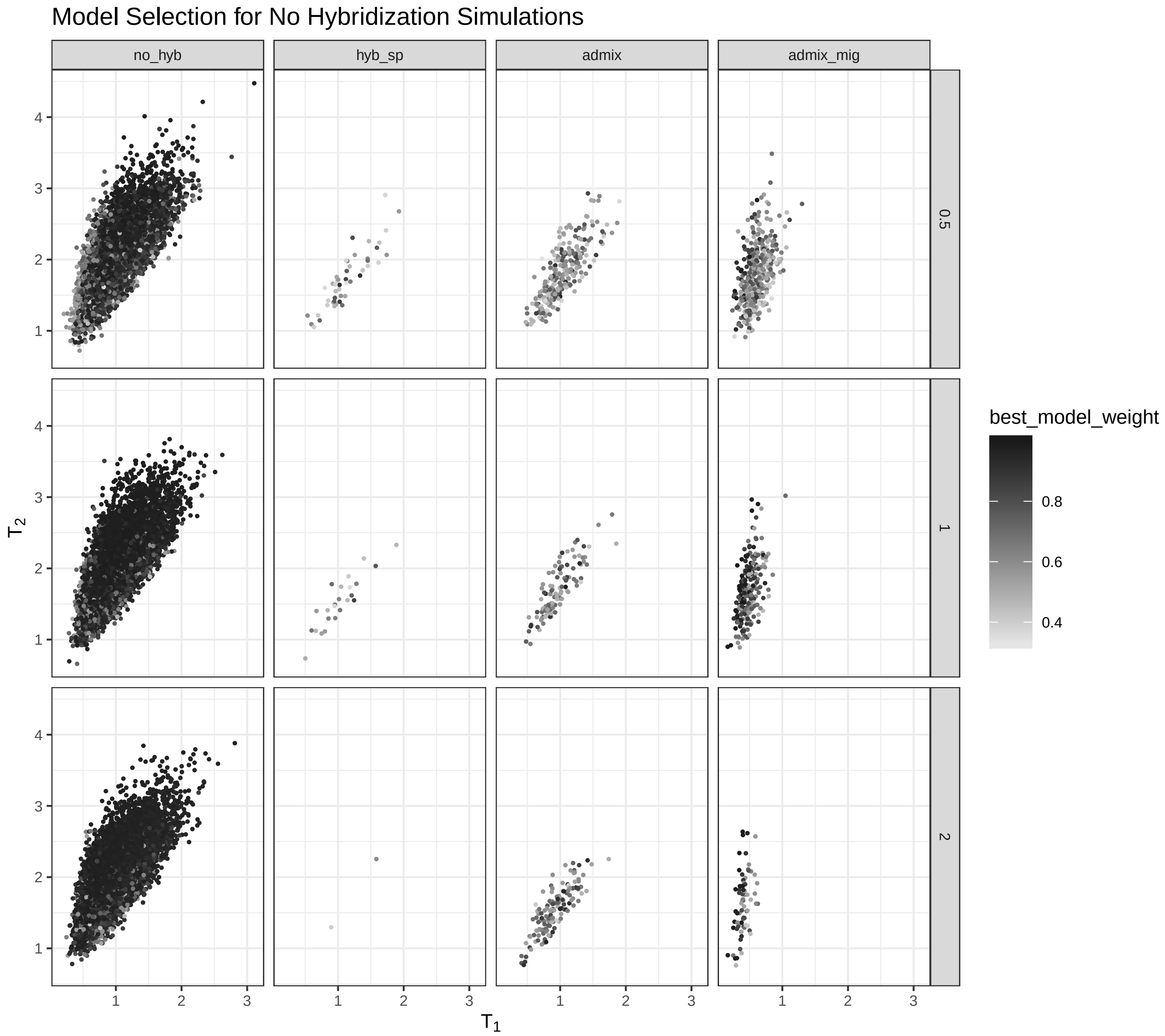

Figure S5: Model prediction for test data simulated under the no hybridization scenario. Columns represent the model chosen by the HyDe-CNN architecture trained on minimum $d_{XY}$ and rows show different branch scalings. The divergence time between $P_1$ and $P_2$ ($T_1$) is plotted on the x-axis and the divergence time between $P_3$ and $(P_1, P_2)$ ($T_2$) is plotted on the y-axis.

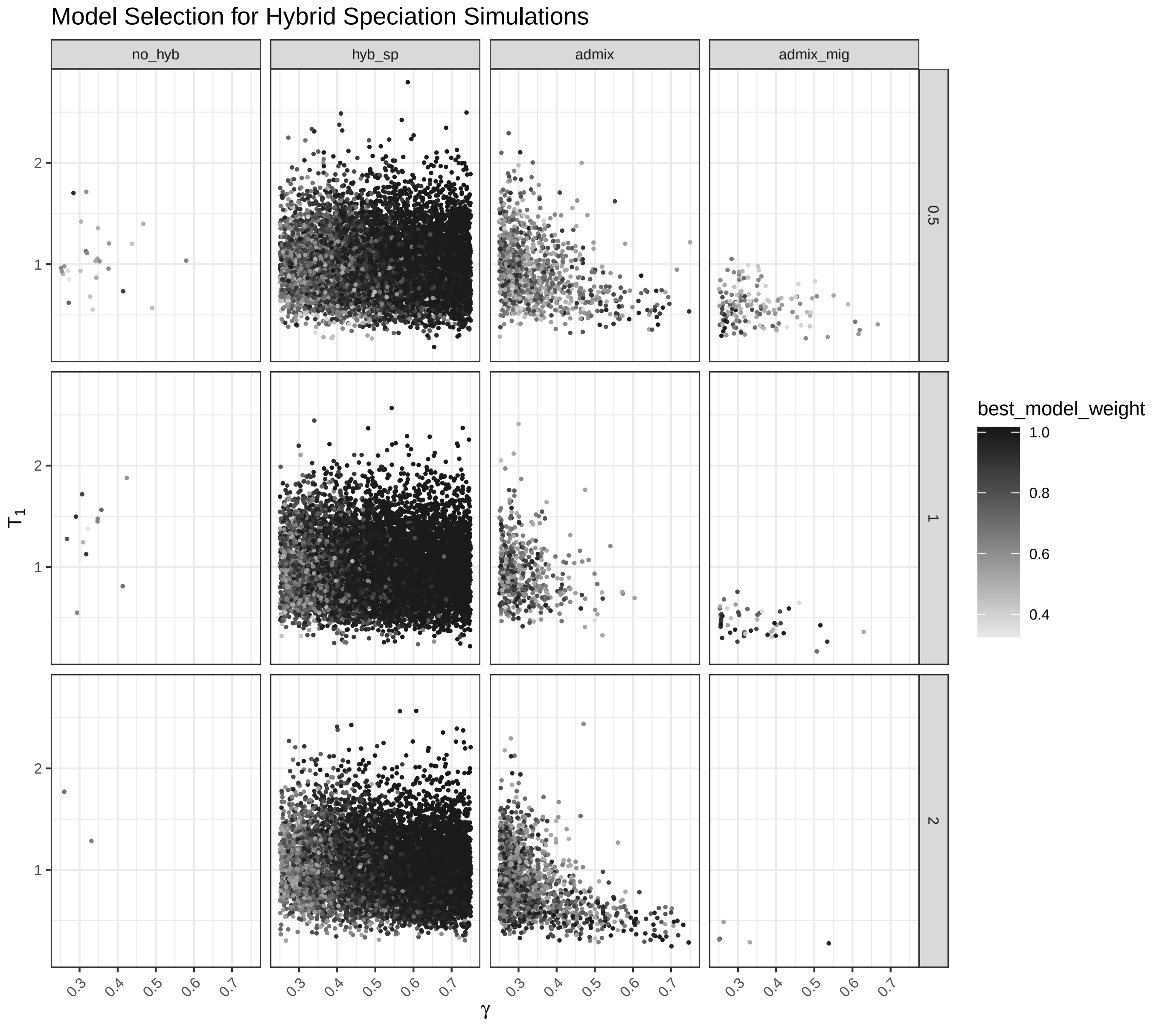

Figure S6: Model prediction for test data simulated under the hybrid speciation scenario. Columns represent the model chosen by the HyDe-CNN architecture trained on minimum $d_{XY}$ and rows show different branch scalings. The hybridization fraction ($\gamma$) is plotted on the x-axis and the divergence time between $P_1$ and $P_2$ ($T_1$) is plotted on the y-axis.

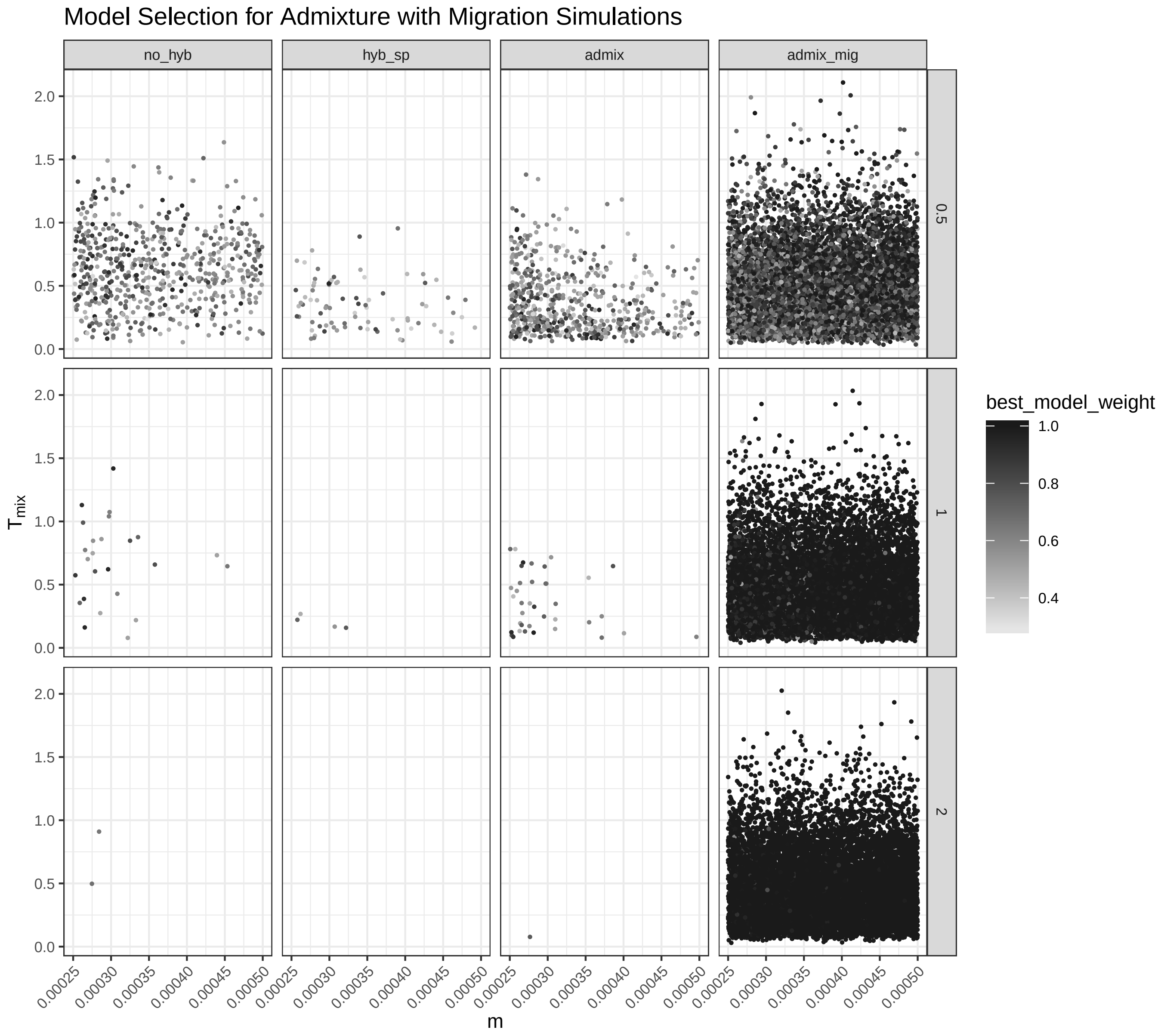

Figure S7: Model prediction for test data simulated under the admixture with gene flow scenario. Columns represent the model chosen by the HyDe-CNN architecture trained on minimum $d_{XY}$ and rows show different branch scalings. The migration rate $m$ is plotted on the x-axis and the timing of admixture ($T_{mix}$) is plotted on the y-axis.

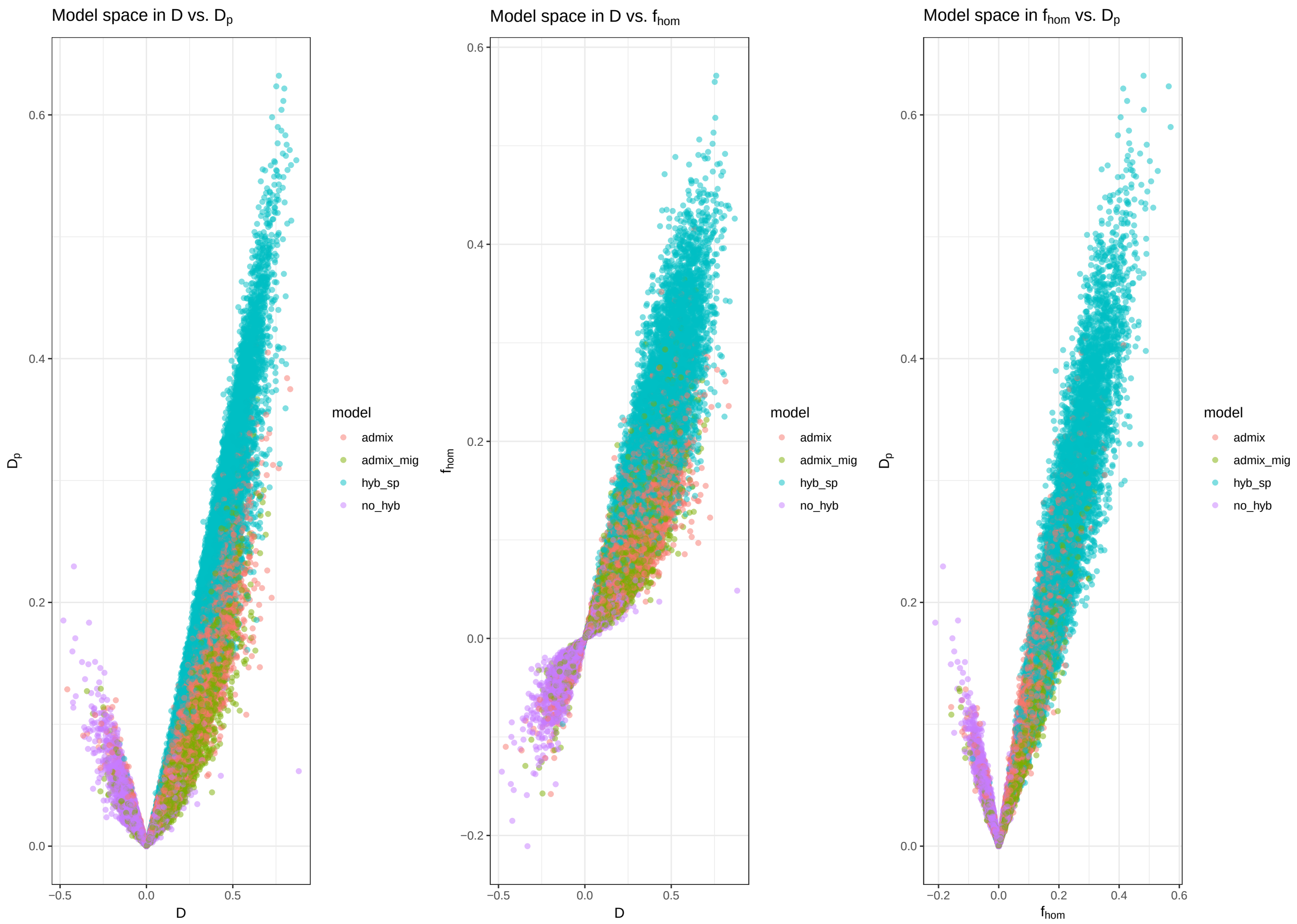

Figure S8: Pairwise plots of calculated summary statistics ($D$, $f_{hom}$, $D_p$) for training a random forest classifier at the 0.5 coalescent unit branch scaling. The color of each dot corresponds with a single simulation and represents its generating model.

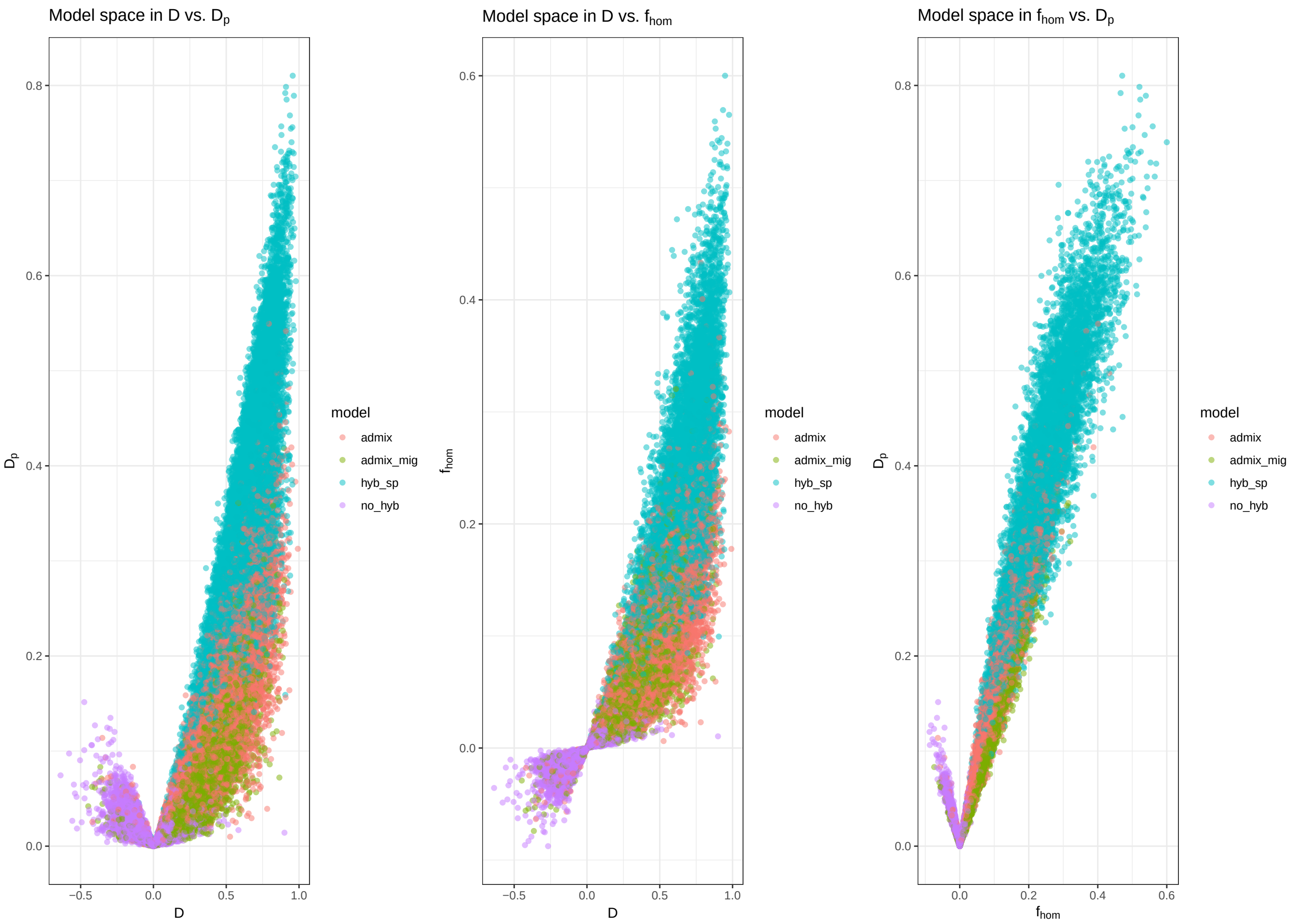

Figure S9: Pairwise plots of calculated summary statistics ($D$, $f_{hom}$, $D_p$) for training a random forest classifier at the 1.0 coalescent unit branch scaling. The color of each dot corresponds with a single simulation and represents its generating model.

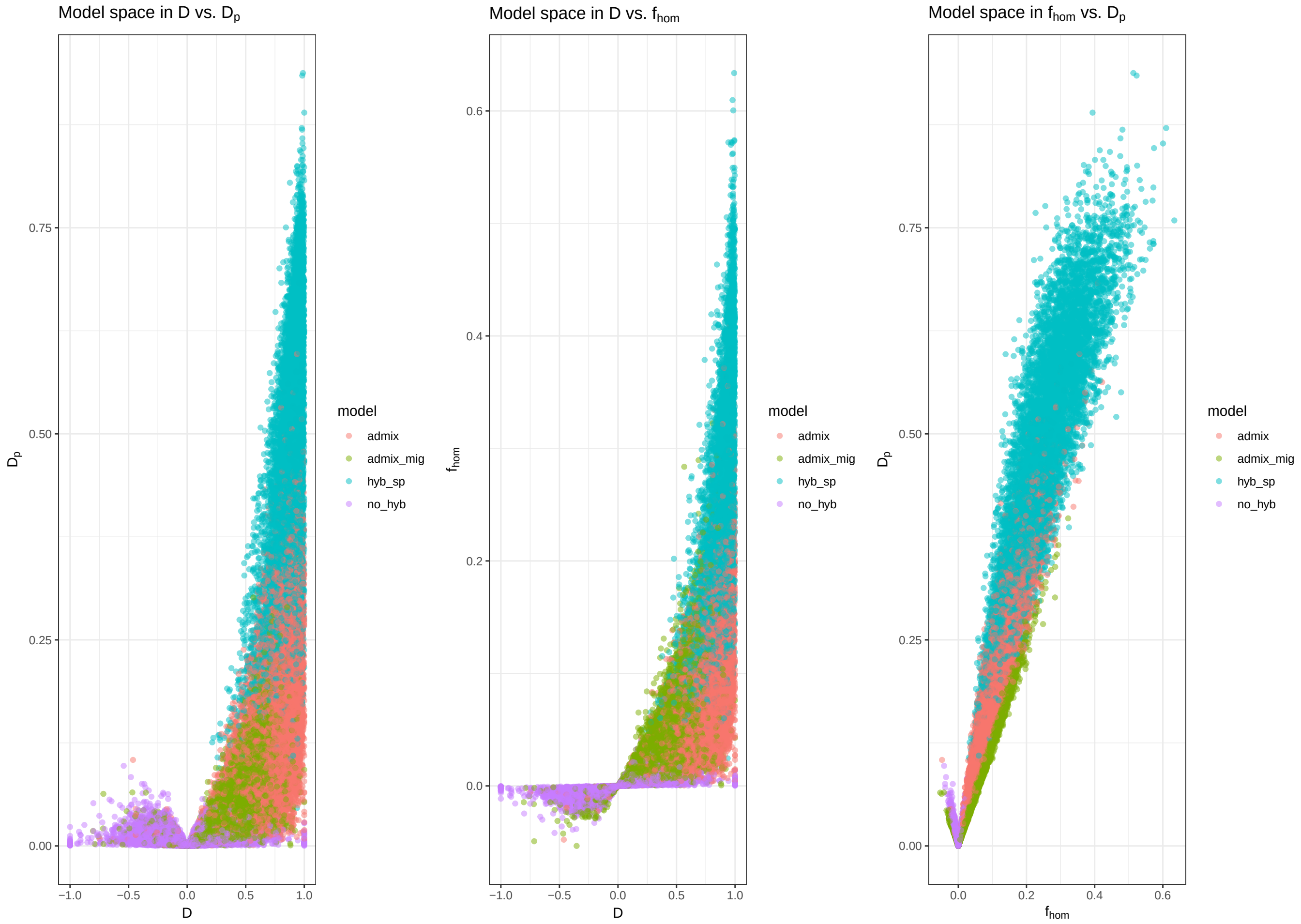

Figure S10: Pairwise plots of calculated summary statistics ($D$, $f_{hom}$, $D_p$) for training a random forest classifier at the 2.0 coalescent unit branch scaling. The color of each dot corresponds with a single simulation and represents its generating model.

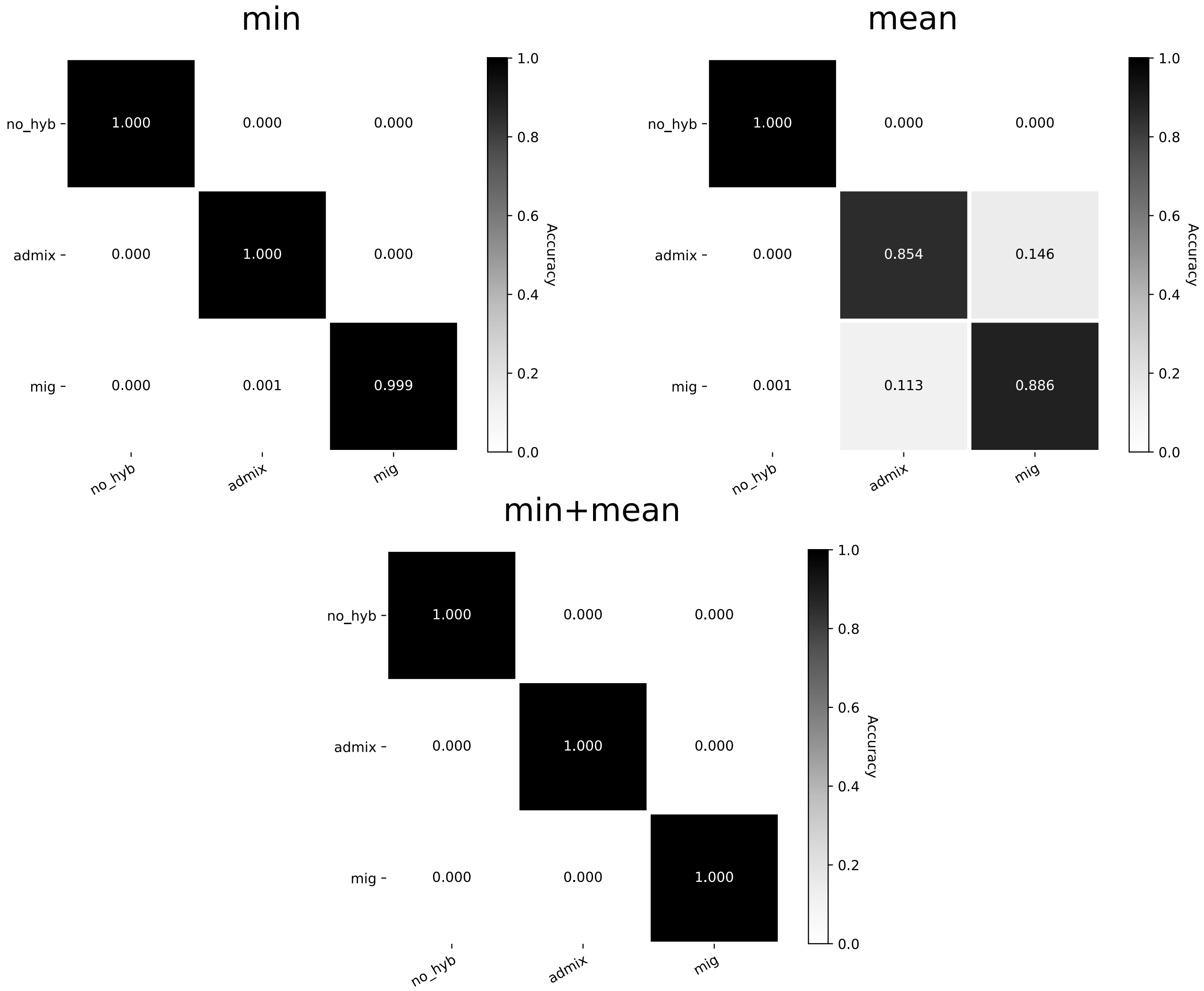

Figure S11: Confusion matrices for images simulated under the demographic models (no hybridization [no_hyb], admixture [admix], and continuous migration [mig]) tested in *Heliconius* across the different input types.
